# Treatment utilization and medical problems in a community sample of adult women with anorexia nervosa

**DOI:** 10.1101/547133

**Authors:** Brooks Brodrick, Jessica A. Harper, Erin Van Enkevort, Carrie J. McAdams

**Author notes:** **Correspondence:** Brooks Brodrick, M.D., Ph.D.

## Abstract

Anorexia nervosa has a prolonged course of illness, making both defining recovery and determining optimal outpatient treatments difficult. Here, we report the types of treatments utilized in a naturalistic sample of adult women with anorexia nervosa in Texas. Participants were recruited from earlier studies of women with anorexia nervosa (AN-C, n = 28) and in weight recovery following anorexia nervosa (AN-WR, n = 18). Participants provided information about both their illness and treatments during their most severe period (severe period) as well as during the two to six years following original assessments (follow-up period). New follow-up groups were defined based on current clinical status (continued eating disorder, AN-CC; newly in recovery, AN-CR; sustained weight-recovery, AN-WS), and clinical utilization was compared across groups. There were no differences in groups related to symptoms or treatments utilized during the severe-period. During the follow-up period, intensive outpatient programs were utilized significantly more by the AN-CC group than the other groups, and dietitians were seen significantly less by the AN-WS group. Medical complications related to the ED were significantly more common in the AN-CC group. All groups maintained similar levels of contact with outpatient psychiatrists, therapists, and primary care physicians.

## 1 Introduction

Anorexia nervosa (AN) is a serious mental illness characterized by difficulties consuming sufficient calories to maintain body weight in conjunction with disturbances in self-perception. Although this disease often begins in adolescents and young adulthood, the course of disease is prolonged and outcomes are poor (1). In examination of treatment outcomes in partial hospitalization and intensive outpatient settings, individuals with AN had worse outcomes than individuals with bulimia nervosa, binge eating disorder, and other specified feeding or eating disorder (2). Additionally, a recent meta-analysis showed that treatment changed only short-term weight outcomes but did not significantly impact either short-term or long-term psychosocial outcomes or long-term recovery (3).

AN has a relapse rate of approximately 31%, with the highest risk in the first year to two years post-treatment (4). The dropout rate from outpatient treatment for AN is estimated to range from 20-40% (5). Qualitative research on recovered women highlights the importance of psychosocial support as part of recovery (6). However, there is little evidence that defines what is optimal outpatient treatment for adults with AN, and minimal data describing even the types of treatment utilized in the community in the United States. Most longitudinal work in adults has focused on assessment of changes in clinical symptoms (1, 7) or follow-up of patients after specific types of interventions (8, 9).

Here, we evaluated the symptoms and types of treatments utilized by women with AN in Texas. All participants were diagnosed with AN previously, and were originally recruited to participate in studies comparing women with AN currently and those in long-term weight recovery following AN. Previously, we reported on the re-assessed clinical status of these participants two to six years after original measures were collected, comparing clinical and cognitive symptoms at baseline and outcome in participants with continued symptoms, participants whose symptoms had remitted since initial assessments, and participants who remained in sustained recovery (7). Here, we report and compare the types of treatment obtained by these subjects at the most severe stage of their eating disorder as well as during the time period since participating in the study. We hypothesized that there would be less utilization of outpatient treatment providers amongst the women that relapsed and persisted with disease relative to those that recovered.

## Methods

### 2.1 Participants

Forty-six women ages 20-61 (*M* = 30.3, *SD* = 8.61) participated. All subjects had completed baseline measures as part of one or more of three previous studies (enrollment period was 2011-2014) including both currently-ill (AN-C) and weight-recovered (AN-WR) women with AN (10-12). For all studies, the AN-C group included women who had met DSM-IV criteria for AN within the previous six months whereas the AN-WR group required women to have maintained a body mass index (BMI) > 19 for at least 12 months despite having met DSM-IV criteria for AN in their lifetime. In addition, the AN-WR participants could not have met DSM-IV criteria for bulimia nervosa in the last two years. The studies were conducted in overlapping time periods; many participants were in more than one study, with the variation resulting from subject preferences and safety concerns related to participation in magnetic resonance imaging, blood draws, and genetic analyses. Participants provided written informed consent approved by the UT Southwestern Institutional Review Board. The Structured Clinical Interview for DSM-IV (13) was conducted upon initial enrollment to confirm current or past AN and comorbid diagnoses; all interviews were conducted by trained assessors at the masters or doctoral level. Follow up self-reports were collected and managed using REDCap electronic data capture tools hosted at UT Southwestern (14), where participants also provided digital consent upon beginning their follow-up surveys.

### 2.2 Outcome criteria

The primary outcome criterion was based on maintaining a BMI > 19 for at least 12 months without meeting criteria for bulimia nervosa; this matched the enrollment criteria utilized for AN-WR in the original studies. Participants in recovery could not have participated in intensive outpatient, partial hospital, residential, or inpatient treatment for at least 12 months. Participants were characterized as still ill (AN-CC), in recovery from a baseline ill status (AN-CR), or in sustained weight recovery (AN-WS).

### 2.3 Measurements of treatment

Using a guided, semi-structured interview, each participant provided information about their eating disorder, focusing on characterization of their symptoms and interactions with clinical treatment providers. With regard to their eating disorder, each subject self-reported their age of onset, their age of first treatment, and their age during their worst period with the eating disorder. Body mass index, all care utilized immediately following the worst period (inpatient/residential, partial hospital, intensive outpatient, or outpatient), and the symptoms (restriction, binge-ing, purging, depression, anxiety, alcohol, and other substance use) during the worst period were detailed. The duration and intensity of outpatient treatment following this period was queried, including interactions with five different types of clinical care (primary care physician, psychiatrist, therapist, dietitian, group therapy). If answered affirmatively that type of clinical care was sought, then the frequency (annually, quarterly, monthly, biweekly, weekly) and duration of that clinical interaction was inquired.

Separately, treatment utilization following the individuals’ participation in the research study was assessed. Two tables were completed for every subsequent six month period to present. The first table assessed a need for a higher level of care for the eating disorder (or a related comorbid condition such as suicidality or substance abuse). All hospitalizations, including both medical and psychiatric (further classified as inpatient, residential, partial hospital, intensive outpatient), and ER visits. The second table assessed the participant’s interaction with five different types of outpatient clinical care, including frequency and duration, for each 6 month period.

### 2.4 Statistical analysis

Statistical analyses were conducted using the IBM Statistical Package for Social Sciences (SPSS; v.23). To determine differences between groups, separate one-way analysis of variance (ANOVAs) were conducted on continuous data. Post hoc pairwise comparisons were conducted using a Bonferroni correction for multiple comparisons. Categorical and dichotomous (yes/no) data were analyzed by Chi-square analyses.

## Results

### 3.1 Participants

As previously reported, amongst the AN-C group, 17 participants remained ill (AN-CC) and 11 achieved recovery from a baseline ill status (AN-CR), a recovery rate of 39%. In addition, all 18 of the AN-WR group remained weight-recovered at follow up (AN-WS). There were no differences in the distributions of subtypes of AN by group (AN binge-purge/restricting, AN-CC 9/8, AN-CR 5/6, AN-WS 11/7; *X^2^*(2) = 0.67, *p* = 0.71, *V* = 0.12).

The AN-CR group had a later age of onset than the AN-WS (age of onset in years, AN-CC, 16.3, AN-CR 19.2, AN-WR 14.2, *F*_(2, 43)_ = 3.86, *p* = 0.03; post-hoc AN-WS < AN-CR *p* = 0.02). The AN-CC group had an earlier age of first treatment than the AN-CR group (age of first treatment, AN-CC 17.8, AN-CR 22.7, AN-WR 18.5, F(2, 42)= 3.35, *p* = 0.045; post-hoc AN-CC < AN-CR, *p* = 0.05).

### 3.2 Severe-period symptoms and treatments

There were no differences in the clinical symptoms or treatments utilized in the worst period of the disease for any of the groups (Table 1). Eating disorder symptoms were considered most severe for all subjects on average at 22.6 years, and the lowest BMI was similar across all subjects (mean lowest BMI 15.32). Across all groups, 43.6% of subjects reported binge-eating behaviors, and 50% reported purging symptoms in the severe period. 84.8% of the subjects had comorbid depression and 82.6% had comorbid anxiety. Problems with alcohol and substance abuse were less prevalent across the sample, with 21.7% of subjects reporting an alcohol use disorder and 10.9% reporting substance abuse. No differences in the types of intensive treatment utilized following that severe-period were observed across the 3 groups, with 43.5% utilizing inpatient or residential treatments, 41.3% in partial hospital programs, 39.1% in intensive outpatient.

**Table 1.** Participant Characteristics

|  | AN-CC<br>(N = 17) | AN-CR<br>(N = 11) | AN-WS<br>(N = 18) | Statistical Comparisons |  |
| --- | --- | --- | --- | --- | --- |
|  | Mean(SD) | Mean(SD) | Mean(SD) | Test Statistic | p-value |
| Age of Onset | 16.29(3.10) | 19.18(7.81) | 14.22(3.21) | $F(2,43) = 3.86$ | **.03** |
| Age of First Treatment | 17.82(3.03)<sup>a</sup> | 22.73(7.92)<sup>a</sup> | 18.53(4.35) | $F(2,42) = 3.35$ | **.045** |
| Baseline BMI | 17.54(1.52)<sup>a</sup> | 18.39(1.08)<sup>b</sup> | 22.87(2.61)<sup>ab</sup> | $F(2,43) = 36.38$ | **<.001** |
| Lowest BMI | 14.59(2.05) | 15.72(1.60) | 15.64(1.59) | $F(2,43) = 1.97$ | .15 |
|  | n(%) | n(%) | n(%) |  |  |
| Race | | | | $X^2(6) = 8.73$ | .19 |
| White | 17(100 %) | 11(100%) | 13(72%) |  |  |
| Asian | 0(0%) | 0(0%) | 2(11%) |  |  |
| Native American | 0(0%) | 0(0%) | 2(11%) |  |  |
| African American | 0(0%) | 0(0%) | 1(6%) |  |  |
| Ethnicity | | | | $X^2(2) = 1.56$ | .46 |
| Hispanic | 2(12%) | 0(0%) | 1(6%) |  |  |
| Non-hispanic | 15(88%) | 11(100%) | 17(94%) |  |  |
| Education (years) | | | | $X^2(4) = 1.93$ | .75 |
| ≤12 | 4(24%) | 1(9%) | 2(11%) |  |  |
| 13-16 | 8(47%) | 6(55%) | 8(44%) |  |  |
| >16 | 5(29%) | 4(36%) | 8(44%) |  |  |

BMI = Body Mass Index; Means with the same superscript are significantly different from one another (Bonferroni correction for multiple comparisons); Significant differences are in bold.

### 3.3 Follow-up period symptoms and treatments

Few differences were observed in the types and amount of care obtained from clinical providers in the follow-up period between initial participation in the study and returning for the follow-up measures (Table 3). Some members in all groups attended inpatient, residential or partial hospital eating disorder programs (AN-CC, 47%, AN-CR, 36%, AN-WS, 17%). The three subjects in the AN-WS group that required a higher level of care did so for escalating ED symptoms consistent with having developed and then recovered from bulimia nervosa during the follow-up period. The only significant difference was related to intensive outpatient treatment programs, which were most heavily utilized by those currently ill with AN (AN-CC, 53%, AN-CR, 9%, AN-WR, 6%, *X^2^*(2) = 12.53, *p* = 0.002).

**Table 2.** Symptom and Treatment Comparisons for Most Severe Period

|  | AN-CC<br>(*N* = 17) | AN-CR<br>(*N* = 11) | AN-WS<br>(*N* = 18) | Statistical Comparisons |  |
| --- | --- | --- | --- | --- | --- |
|  | Mean(SD) | Mean(SD) | Mean(SD) | Test Statistic | *p*-value |
| Age at Enrollment | 25.00(6.85) | 26.00(6.39) | 29.44(10.37) | $F(2,43) = 1.33$ | .27 |
| Age, Most Severe Period | 22.97(8.32) | 24.18(6.94) | 20.65(7.33) | $F(2,42) = 0.79$ | .46 |
| Lowest BMI | 14.59(2.05) | 15.72(1.60) | 15.64(1.59) | $F(2,43) = 1.97$ | .15 |
|  | n(%) | n(%) | n(%) |  |  |
| Symptoms |  |  |  |  |  |
| Restricting | 17(100%) | 11(100%) | 18(100%) |  |  |
| Binging | 8(47%) | 4(36%) | 8(44%) | $\chi^2(2) = .32$ | .85 |
| Purging | 11(65 %) | 6(55%) | 6(33%) | $\chi^2(2) = 3.56$ | .17 |
| Depression (Lifetime) | 15(88%) | 8(73%) | 16(89%) | $\chi^2(2) = 1.63$ | .44 |
| Anxiety (Lifetime) | 14(82%) | 10(91%) | 14(78%) | $\chi^2(2) = .82$ | .66 |
| Alcohol Use (Lifetime) | 3(18%) | 2(18%) | 5(28%) | $\chi^2(2) = .64$ | .73 |
| Other Substance Use (Lifetime) | 2(12%) | 1(9%) | 2(11%) | $\chi^2(2) = .05$ | .98 |
| Treatment |  |  |  |  |  |
| Inpatient | 10(59%) | 4(36%) | 6(33%) | $\chi^2(2) = 2.61$ | .27 |
| Partial Hospitalization | 9(53%) | 6(55%) | 4(22%) | $\chi^2(2) = 4.45$ | .11 |
| Intensive Outpatient | 10(59%) | 4(36%) | 4(22%) | $\chi^2(2) = 4.96$ | .08 |

**Table 3.**
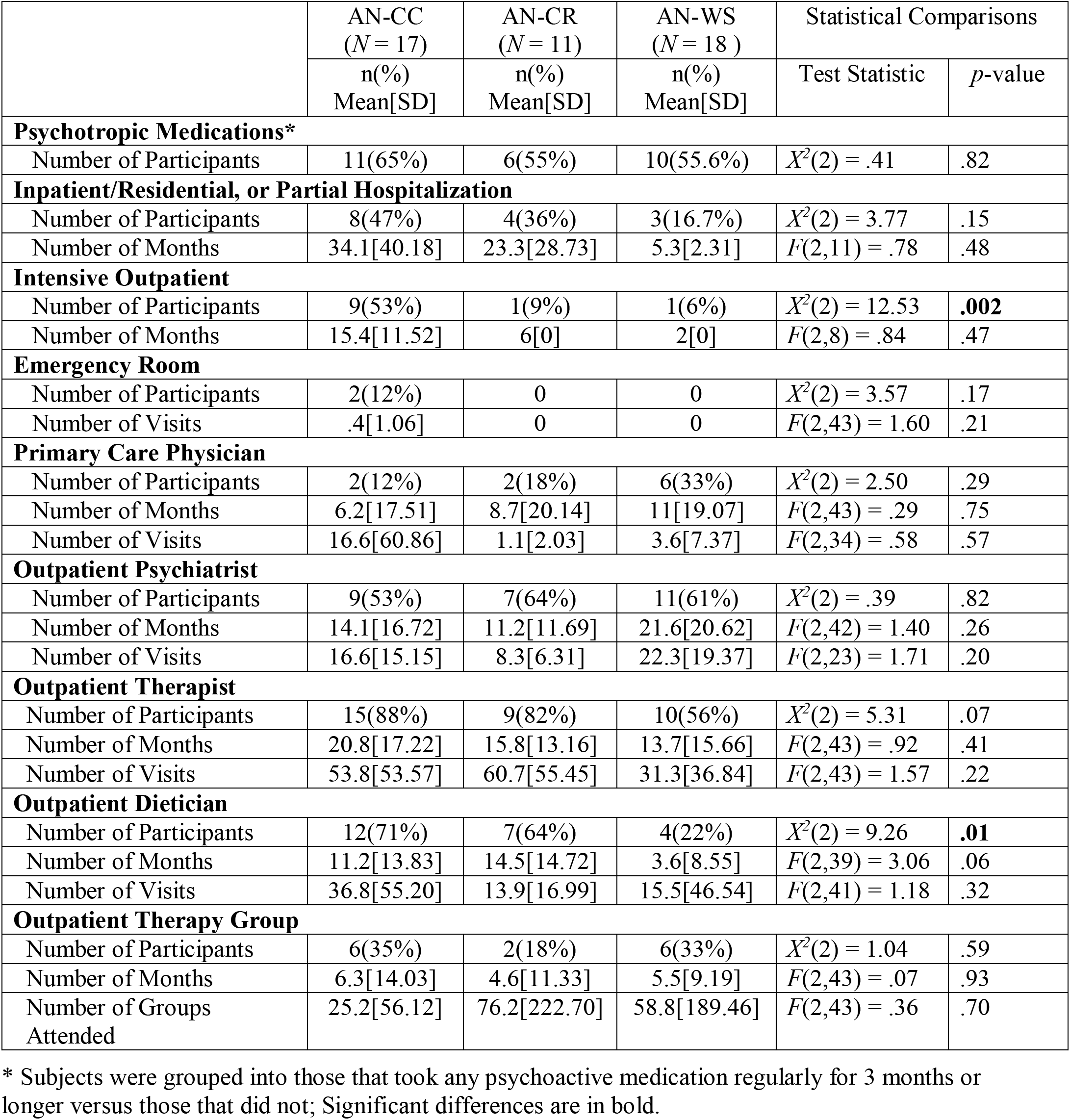
Treatments in Follow-Up Period

In terms of outpatient treatment, similar engagement in outpatient care was observed across all groups. No differences were observed in terms of contact with type of provider across groups with 58.7% in treatment with an outpatient psychiatrist, 73.9% reporting treatment with an outpatient therapist, and 30.4% engaging in group therapy. The more recently ill participants were more likely to see dietitians than those in sustained weight-recovery (AN-CC, 70%, AN-CR 64%, AN-WR 22%, *X^2^*(2) = 9.26, *p* = 0.01).

### 3.4 Medications and medical complications

In this retrospective study, virtually no participants were able to recall either the duration or names of all psychoactive medications taken. Many reported using medications while in inpatient/residential treatment programs but stopping them days or weeks after discharge for a wide variety of reasons (forgot to pick up, did not have a refill, did not think it was helpful). Because of this limited recall, subjects were grouped into those that took any psychoactive medication regularly for 3 months or longer versus those that did not. All participants that took a medication for more than three months appeared confident about their behavior. With these limits, no differences in medication use were seen across the three groups (Table 3).

Participants were also queried about the presence of any other medical problems and their awareness of any medical illnesses or complications derived from the eating disorder (osteopenia/osteoporosis, anemia, electrolyte disturbances, cardiovascular problems) or likely to impact eating behaviors (gastrointestinal disorders). The AN-CC group reported significantly more ED-related medical issues (osteopenia/osteoporosis (5), hypoglycemia, anemia (2), hypophosphatemia) than the AN-CR (inflammatory bowel disease) and AN-WS (osteopenia, irritable bowel) groups (ED medical complications, AN-CC, 52.9%, AN-CR, 9.1%, AN-WR, 16.7%, *χ*^2^ (2) = 8.29, *p* = 0.02). There were not significant differences by groups for other medical problems (non-ED medical problems, AN-CC, 0%, AN-CR, 18.2%, AN-WR, 27.8%, *χ*^2^(2) = 5.33, *p* = 0.07). The non-ED medical problems reported included migraines (2), fibromyalgia, thyroid disorder (2), rheumatoid arthritis, osteoarthritis, pelvic pain, and asthma. There were no differences in engagement with a primary care provider across the groups (percent with primary care provider, AN-CC, 11.8%, AN-CR, 18.2%, AN-WR, 33.3%, *χ*^2^(2) = 2.50, *p* = 0.29).

## 4 Discussion

This work characterizes both the medical and psychiatric symptoms and types of care accessed by a community sample of women living with and recovered from AN in the United States. A first and necessary step towards improving clinical outcomes for AN as it enables us to consider whether the impediments to recovery from AN are related primarily to access to care or the effectiveness of the care that is delivered. Most importantly, we found little evidence that the type of treatments utilized, either at the most severe stage of disease or in outpatient care, were related to longer-term clinical outcomes. Further, we found that most participants were engaged in regular outpatient treatment with therapists, dietitians, and psychiatrists. Another key finding is that the utilization of therapists and psychiatrists did not differ even for the women with years of recovery from AN relative to those more recently ill, suggesting that continued engagement with mental health professionals may be important for all patients with a history of AN.

Failure in treatment of AN is commonly ascribed to the egosyntonic nature of the disorder, with the resulting perception that women with the illness are deliberately choosing not to recover and are at fault (15). However, the reality of the prolonged disease is that it gradually becomes ego-dystonic, as the patients recognize that they have actually lost the ability to eat sufficient food for their mental and physical health (16), and express ambivalence towards the disease (17). Nevertheless, there remains a perspective that patients with AN choose to maintain their illness and could recover if able to utilize already-existing interventions (18-20). This longitudinal data, albeit from a small sample, suggests otherwise. Somewhat surprisingly, all three groups of subjects – defined by their long-term clinical outcomes – engaged in the same types of treatments, with the only exception being more intensive outpatient treatment for those that continued with disease and increased treatment with outpatient dietitians in those more recently ill. Thus, ED treatment utilization was not correlated positively with sustained weight-recovery from AN. Different approaches to treating this disease are desperately needed in the United States. The medical profession should recognize that current treatments for AN are ineffective for many patients, and this should not be viewed as the patients’ choosing to stay sick any more than a lack of response of one’s cancer to chemotherapy is a choice that those patients make.

Clinical symptoms commonly tracked in eating disorders such as severity of disease based on body mass index, presence of bingeing or purging behaviors, and comorbid anxiety or depressive disorders were also unrelated to long-term course of illness. Most subjects experienced depression and anxiety during their eating disorder. Binge-eating behaviors and purging symptoms were observed in about half of the participants, irrespective of group. These data are consistent with recent literature that suggests severity-specifiers for AN may not be clinically relevant (21, 22). In a recent meta-analysis of treatments for AN, Murray et al. (3) found that although weight improved amongst AN participants between beginning and end of treatment, weight was not improved at follow-up, and psychological changes were not observed at either time point. We previously reported that this cohort had no changes longitudinally in body shape questionnaire, anxiety, or self-esteem, although depression and externalizing bias did change over time across groups (7). Another novel finding is that all groups of patients reported that the most severe time-period for eating disorder symptomatology was in their early twenties, potentially identifying a specific high-risk time period of emerging adulthood that could be targeted by increased psychosocial supports or clinical monitoring for eating disorder symptoms.

Another potential gap in the treatment of AN identified here is in the recognition and treatment of the many medical complications and problems related to development or progression of the disorder. More of the AN-CC group reported significant medical complications from their disease than those in both the AN-CR and AN-WR groups. These medical complications included gastrointestinal disturbances (gastroesophageal reflux disorder, irritable bowel disorder, gastroparesis) as well as problems from chronic malnutrition (osteopenia, anemia) and purging behaviors (orthostatic hypotension, electrolyte abnormalities). The presence of medical complications of AN may provide a better severity specifier more closely related to course of illness than psychiatric symptoms or body mass index alone, but few researchers examining the progress of eating disorder patients include assessments of medical health and comorbidities. Individuals with medical complications of the disease might also be less responsive to existing treatments for AN and related psychiatric comorbidities. It is particularly concerning that despite the increased presence of medical complications, the AN-CC group was no more likely to have a primary care provider than the other two groups. This may be related to the growing difficulty in securing a primary care provider in the United States (23). A multidisciplinary team that includes a primary care provider cognizant of the medical complications and their treatment may be of particular benefit.

This study has many limitations. This was a small sample of individuals with AN, and treatment utilization was based on retrospective report of the patient. Because attending treatment programs often results in substantial disruptions of one’s life, recall related to the higher levels of care may be more accurate than the duration one worked with an outpatient therapist, psychiatrist or dietitian. External documentation, exam, or lab work to assess the presence or absence of medical complications for all participants was not possible. The study was based on DSM-IV criteria for AN rather than DSM-5 as the studies began before DSM-5 was published; given that the stringency for AN was actually reduced in DSM-5, all subjects meet those criteria as well. Clinical, cognitive, neural and genetic data from these subjects has been previously published, but information about their severe periods of illness and clinical utilization has not been reported previously. Larger studies that prospectively monitor AN overtime including evaluation of medical complications of the disease in concert with clinical outcome are needed.

## 6 Author Contributions

BB and CJM designed the study. BB, JH, and CJM collected the data. All authors processed and analyzed data. BB, JH, and CJM contributed to writing and editing the manuscript.

